# Split-Belt walking induces changes in active, but not passive, perception of step length

**DOI:** 10.1101/616631

**Authors:** Carly Sombric, Marcela Gonzalez-Rubio, Gelsy Torres-Oviedo

## Abstract

The estimation of limbs’ position is critical for motor control. While motor adaptation changes the estimation of limb position in volitional arm movements, this has not been observed in locomotion. We hypothesized that split-belt walking with the legs moving at different speeds changes the estimation of the legs’ position when taking a step. Thus, we assessed young subjects’ perception of step length (i.e., inter-feet distance at foot landing) when they moved their legs (active perception) or these were moved by the experimenter (passive perception). Step length’s active, but not passive perception was altered by split-belt walking; indicating that adapted efferent inputs changed the perceived limbs’ position without changing information from sensory signals. These perceptual shifts were sensitive to how they were tested: they were observed in the trailing, but not the leading leg, and they were more salient when tested with short than long step lengths. Our results suggest that sensory changes following motor adaptation might arise from mismatched limb position estimates from different sensory sources (i.e., proprioception and vision), which is less prominent in walking. We also speculate that split-belt walking could improve the deficient perception of step length post-stroke, which contributes to their gait asymmetry impairing patients’ mobility.

## Introduction

Successful motor control requires accurate estimation of limb position, which is achieved through a combination of afferent and efferent signals. Afferent information from sensory signals is used to estimate limb position. In addition, efferent information also contributes to the estimation of limb position through prediction of sensory consequences prior to movement execution. The integration of actual and predicted sensory information, from afferent and efferent signals (respectively), is beneficial because it results in more accurate limb’s estimation in space (e.g., ^1–4^). It has been shown that subjects’ estimation of limb position is altered following motor adaptation to a persistent stimulus (e.g., a constant force ^5^). These adjustments to the estimation of limb position can be due to either adaptation of afferent or efferent information. Shifts in sensory information after motor adaptation have been observed with passive perception tasks during which subjects use afferent information to localize their limb after it has been passively moved (e.g., ^5–12^). Moreover, predicted sensory consequences (from efferent information) rely on internal models that are recalibrated during motor adaptation. Thus, the integration of adapted sensory predictions with actual sensory information leads to changes in limb estimation that are assessed through active perceptual tasks, which require subjects to localize their limb after active movements ^5,10,11,13–15^. In summary, the critical ability to estimate the position of our limbs in space using afferent and efferent information is altered by motor adaptation.

Interestingly, the effects of motor adaptation on the estimation of limb position in walking remains elusive. Namely, a recent study probing the perception of one leg position with respect to the body did not find passive or active perceptual after-effects despite robust motor after-effects ^16^. This might be due to how perceptual after-effects were tested given evidence that passive and active perceptual changes post-adaptation are sensitive to the condition in which they are evaluated ^17–19^. For example, passive perceptual after-effects depend on the limb that is tested ^13^, raising the possibility that clearer perceptual changes could be observed when probing both limbs contributing to each step length. Similarly, active perceptual after-effects might be regulated by test condition such as walking speed; given that this factor alters the magnitude of motor after-effects ^20^, and hence the recalibration of internal models underlying active perceptual shifts. It is important to understand the influence of sensorimotor adaptation on perception of limb position because stroke survivors have difficulty perceiving that their step lengths are asymmetric ^21^. Thus, it is of clinical interest to determine the extent to which the estimation of limb position could be shifted following sensorimotor adaptation in locomotion.

In summary, we investigated the effects of sensorimotor adaptation on the estimation of limb position in walking. We hypothesized that sensorimotor adaptation alters the estimation of step length, which requires concurrent localization of each leg’s position. We further hypothesized that walking condition, such as walking speed and/or step size, would regulate the shifts in active perception of limb positions since walking speed alters motor after-effects ^20^ and walking speed also alters step size (e.g., ^22,23^). To test these hypotheses, we separately investigated shifts in passive and active perception of step length following sensorimotor adaptation in walking. Passive perception was assessed by externally moving subjects’ legs, whereas active perception was evaluated by instructing subjects to take steps of different lengths. Importantly, the active perception task was performed under distinct walking speeds and step sizes to test our hypothesis that active perceptual effects depend on the condition in which they are tested. We assessed both passive and active perception since it is possible that sensorimotor adaptation only induces changes in the integration of afferent and efferent information, but not in afferent information alone.

## Results

### Passive Perception of Step Length was Not Adapted Following Split-belt Walking

We aimed to determine the extent to which locomotor adaptation induced a change in the estimation of limb position purely based on afferent information. To this end, adaptation effects in passive step length perception (i.e., passive perceptual after-effects) and movements (i.e., motor after-effects) were assessed in a group of unimpaired young subjects (n=8, 2 females, 24.8±4.8 yrs.) following a split-belt adaptation paradigm known to induce sensorimotor recalibration (e.g., ^24^). We found motor, but not perceptual after-effects. After-effects were defined as differences in step length (Figure 1a-b) or step length asymmetry (Figure 1c) before and after the Adaptation epoch. These after-effects were significantly different between the split and tied sessions in the motor domain (Figure 1b: paired t-tests: slow leg’s step length: p<0.001; fast leg’s step length: p<0.001; Figure 1c: step length asymmetry: p<0.001). In contrast, we did not observe perceptual after-effects based on sensory information alone. Subjects’ passive perception of their step length was assessed with a task in which participants indicated their estimated step length after the legs were moved by the experimenter to different step length sizes (Figure 1a: short, comfortable, long). We did not observe significant changes in subjects’ perception of the slow leg’s step length across testing sessions (Figure 1a: one-way repeated measures ANOVA testing the effect of session; p=0.98). While we found a significant session effect on the perception of fast leg’s step length (Figure 1a: p=0.035), this constituted a very small perceptual shift (0.8 ± 1.5 cm) compared to the significant motor after-effects in the fast leg’s step length (11.7 ± 5.9 cm) and the slow leg’s step length (−20.1 ± 6.7 cm). Thus, we observed a significant, but marginal adaptation effect on the perception of the fast leg’s step length. In sum, split-belt walking produced robust motor after-effects, but did not change subjects’ estimation of step lengths based on sensory (afferent) information alone.

**Figure 1.**
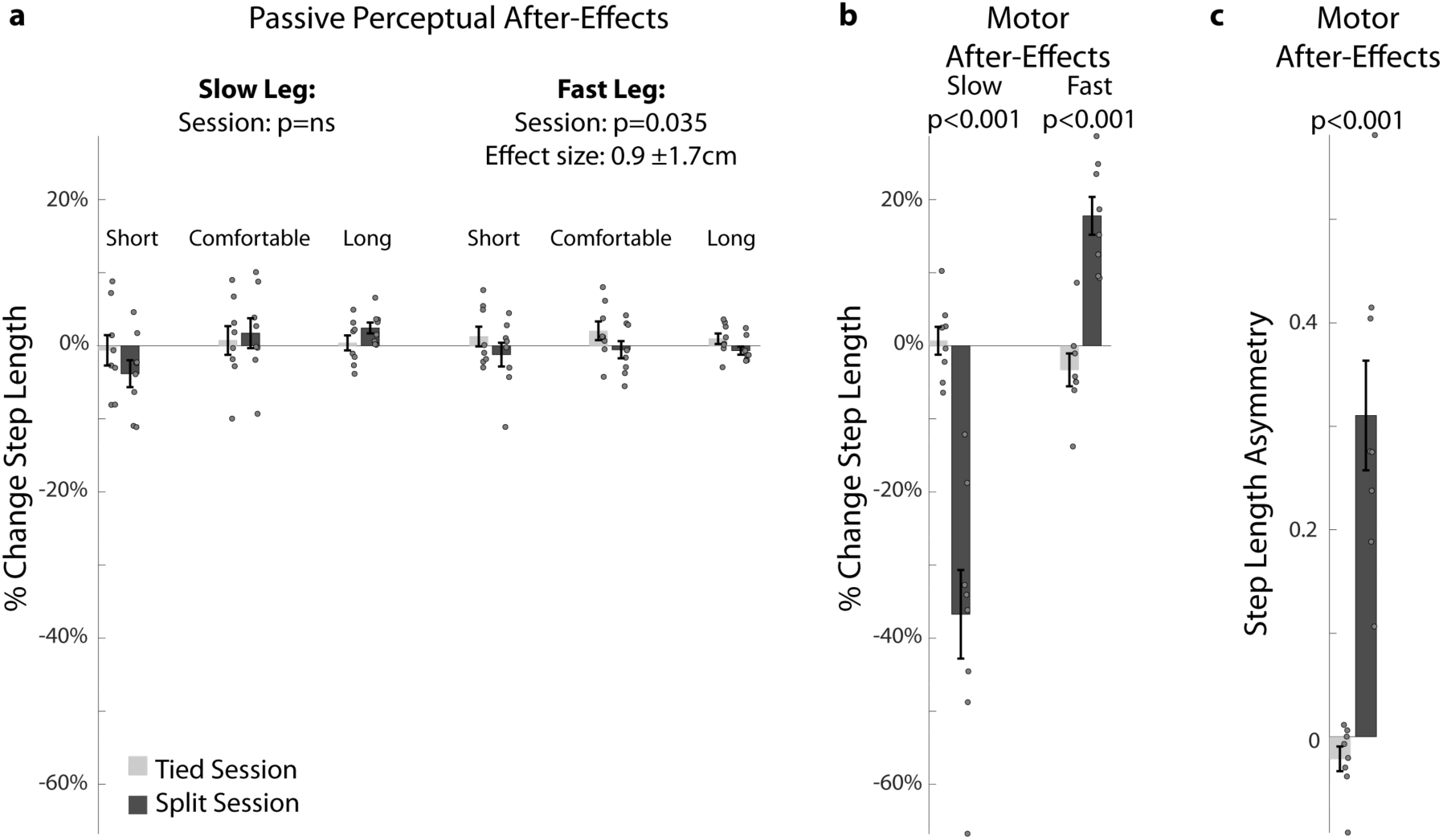
Split-belt walking did not induce changes in passive perception of step lengths. **(a-b)** In all plots, step length is defined as the distance between the ankles at heel strike (i.e., when the foot hits the ground). Change in step length was computed as the difference in step length before and after the Adaptation epoch, which during the control session was an extended period of tied walking (light grey) and during the split session was an extended period of split walking (dark grey). These changes were expressed as a percentage of each subject’s mean step length during baseline walking at 1 m/s to account for different step sizes across individuals. Negative and positive values indicate that step lengths were either perceived shorter or longer, respectively, after the Adaptation epoch compared to before. **(a) Passive Perception After-Effects**: Bars’ height indicates group mean changes in perception after the Adaptation epoch relative to baseline behavior. Dots indicate values for each individual and error bars indicate standard errors. Passive perception was quantified by probing subjects’ perception of three step length sizes (short, comfortable, and long) imposed by passively moving subjects’ legs. Each participant’s weight was equally distributed between the legs when their step length perception was probed. **(b) Motor After-Effects of individual legs:** Bars’ height indicates group mean changes in step length after either the tied session or the split session. Standard errors and individual values (dots) are also displayed. **(c) Motor After-Effects of step length asymmetry**: Step length asymmetry is defined as the differences between step lengths of two consecutive steps normalized by the sum of these step lengths. Positive values indicate that steps taken with the dominant leg are longer compared to those taken with the non-dominant leg; and vice versa for negative values. Bar height indicates group mean changes in step length symmetry after the Adaptation epoch. Standard errors and individual values are also displayed.

### Active Perception of Step Length was Modulated in Both Legs After Split-Belt Walking

Unlike passive perception, subjects’ active perception of step length based on efferent information changed after split-belt walking compared to after tied walking. We evaluated subjects’ (n=27, 16 females, 25.1±5.4 yrs.) active perception of step length by prompting step lengths of two distinct magnitudes (i.e., long or short) on a VR headset (see Methods section). We specifically assessed whether participants overshot or undershot these target step lengths after an Adaptation epoch, which consisted of an extended period of tied walking during the tied session and split-belt walking during the split session. Active perception after-effects were quantified as differences in executed step lengths before and after the Adaptation epoch (i.e., %Change in Step Length). Perceptual after-effects were subsequently compared between the two testing sessions to determine if split-belt adaptation induced changes in the active perception of step length. Importantly, perceptual and motor after-effects were tested under distinct walking speeds (i.e., mid and slow) to determine if walking condition had a similar effect on active perceptual after-effects, as it does on motor after-effects ^20^. Thus, we had a total of three groups labeled by the step length size that was prompted first in the perceptual task and the speed at which it was tested (Long-MidSpeed, Short-MidSpeed, Short-SlowSpeed). Figure 2a shows that following split-belt walking, all groups undershot (negative values) or overshot (positive values) target step lengths with the leg that walked slow (slow leg) or fast (fast leg), respectively, during the Adaptation epoch. This indicated that subjects perceived their step lengths to be either longer or shorter than actual when stepping with the slow or fast leg, respectively. Consistently, we found a significant effect of session on the active perception of the slow (p<0.001) and fast step lengths (p<0.001) with one-way repeated measures ANOVAs. These shifts in perception of individual leg’s step length led to significant differences in step length asymmetries during the active perceptual task across sessions (Figure 2c: session effect: p<0.001).

**Figure 2.**
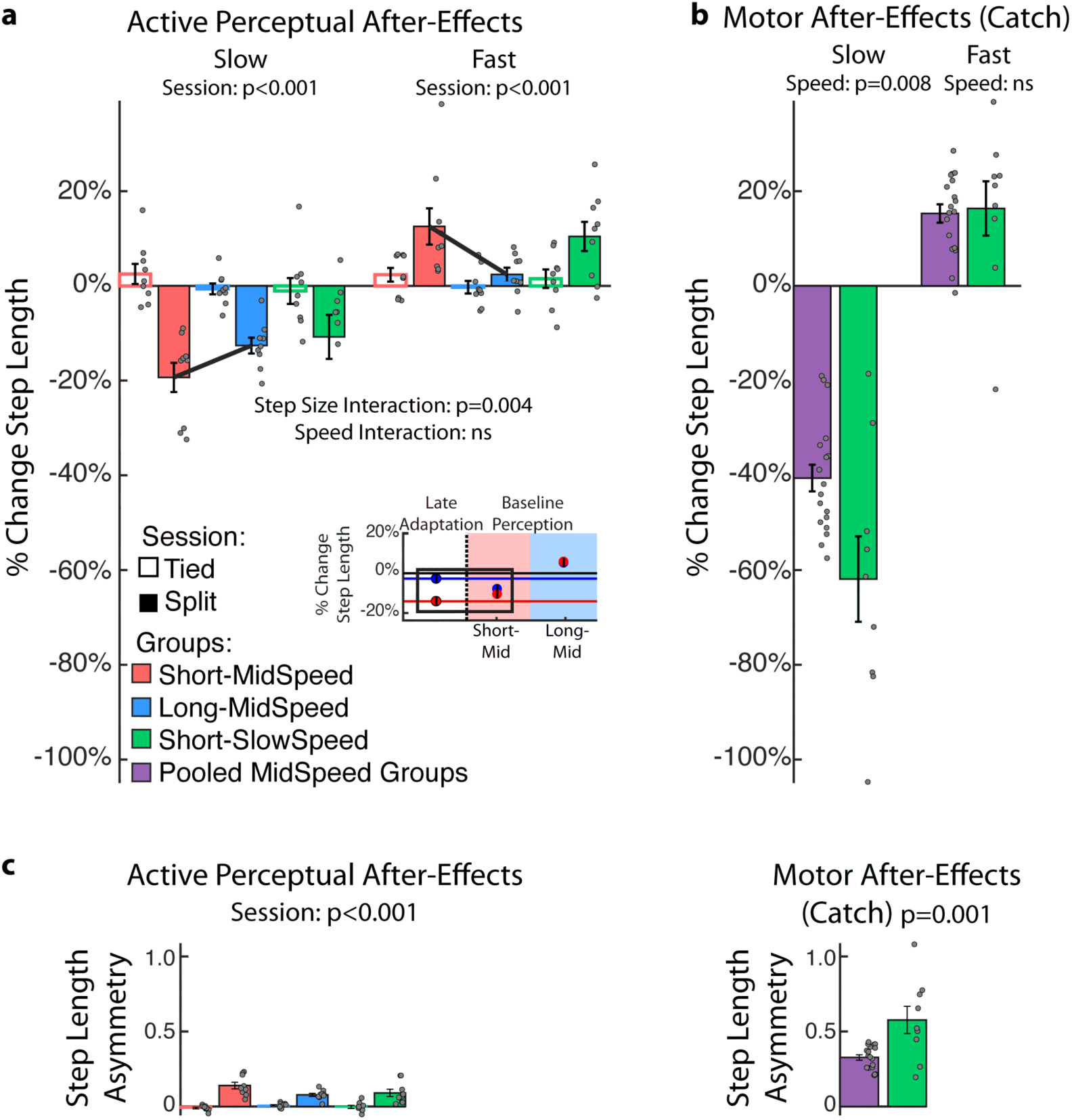
Split-belt walking induces changes in active perception. **(a) Step Length Active Perception After-Effects:** Bars’ height indicates group mean changes in step length before and after the Adaptation epoch, which consisted of tied (empty bars) or split walking (filled bars). These changes were expressed as a percentage of each subject’s mean step length during the baseline active perceptual task tested at the specific step length size and walking speed for each group. For example, we used the mean step lengths during the baseline perceptual task aiming at short step lengths and walking at slow speed for the Short-SlowSpeed group. Negative or positive values indicate that subjects either undershot or overshot the target step lengths, respectively because they perceived step lengths to be longer or shorter than the actual value. The insert illustrates the %Change in Step Length that was executed by the slow (red dot) and fast (blue dot) step lengths at steady state during split-belt walking (Late Adaptation; white background). We also show the target step lengths (red and blue dots) for probing active perception during baseline in the Short-MidSpeed group (red background) and the Long-MidSpeed group (blue background). Standard errors are displayed in all plots. **(b) Step Length Motor After-Effects:** Bars’ height indicates group mean changes in step length relative to baseline walking during the Catch trial tested at either 1.0m/s (Short-MidSpeed and Long-MidSpeed) or 0.5 m/s (Short-SlowSpeed). The mid speed groups were pooled for this analysis since these two groups experienced exactly the same protocol prior to the catch trial. **(c) Step Length Asymmetry Perceptual and Motor After-Effects:** Bars’ height indicates group mean changes in step length asymmetry relative to baseline walking during the Perceptual trials and during the Catch trials. The left panel illustrates the mean step length asymmetry that results from the change in individual step lengths during the active perceptual task for each group. The right panel illustrates the mean step length asymmetry that results from changes during the Catch trial tested at the Mid (1.0 m/s) or Slow (0.5 m/s) walking speed. Standard errors are displayed in all plots.

### Active Perception of Step Length was Modulated by Step Size, but Not Walking Speed

Subsequent analysis on the split session indicated that step length size, but not walking speed, had an effect on active perception after-effects. Specifically, a two-way repeated measures ANOVA indicated a significant interaction between leg and step length size (leg#step length size: *p* =0.004) for the groups starting the perceptual task with different step lengths sizes, but the same walking speed (i.e., Short-MidSpeed vs. Long-MidSpeed). This interaction was driven by greater after-effects in the perception of the fast leg’s step length when taking short (Figure 2a, red bar) compared to long (Figure 2a, blue bar) step lengths (two sample t-test: p=0.024). This might be due to greater similarities between training and testing in the short vs. long conditions. Namely, the executed step lengths during the training condition (i.e., split-belt walking, Late Adaptation in Figure 2a insert, white background) were qualitatively more similar to the target step lengths in the Short-MidSpeeds group (Figure 2a insert, red background) than in the Long-MidSpeeds group (Figure 2a insert, blue background). Conversely, we did not find an interaction between leg and walking speed (leg#speed: *p* =0.16) for the groups that walked at different speeds, but same step lengths in the active perceptual task (i.e., Short-MidSpeed vs. Short-SlowSpeed). This contrasted the effect of walking speed on motor after-effects (Figure 2b). Specifically, the slow step length motor after-effects were larger when tested at slow than mid walking speeds (Figure 2b: two sample t-test between pooled mid speed groups vs. a slow speed group; p=0.008), whereas they were the same for fast step lengths (p=0.83). Thus, the effect of speed on step length asymmetry (Figure 2b: p=0.001) was due to the slow step length after-effects. In summary, split-belt walking induces motor and active perceptual after-effects that are regulated by the way they are tested: motor after-effects are altered by walking speed, whereas perceptual after-effects are altered by step length size.

### Active Perception of Step Length Decayed Without Fully Washing Out Motor After-Effects

We observed that the active perceptual after-effects decayed as they were tested, but their washout did not fully eliminate motor after-effects. Namely, we found that perceptual after-effects decayed exponentially as people walked in the active perceptual task (Figure 3a: *y* = *c* + *ae*^(−*x*/τ)^). This exponential decay was clearly observed in the active perception of the slow step length in all groups (Figure 3a, top panel: τ [95% CI]; Short-MidSpeed: 24.8 [31.0, 20.8], Long-MidSpeed: 17.7 [14.1, 23.6], Short-SlowSpeed: 27.1 [19.4, 45.0]), but only in the fast step length for groups tested with the short target first (Figure 3a, bottom panel: τ [95% CI]; Short-MidSpeed: 25.7 [19.4, 37.8], Long-MidSpeed: 2.4 [-0.86, 3.0], Short-SlowSpeed: 53.2 [37.2, 93.1]). These findings further support that subjects did not have perceptual shifts of the fast leg’s step length when probed with long steps. Interestingly, the active perceptual task did not fully washout motor after-effects. This was tested by comparing perceptual after-effects at the end of the active perceptual task (Figure 3b: Late Perception) and the remaining motor after-effects tested immediately afterwards (Figure 3b: Early Post-Adaptation). Of note, the belts moved at the same tied speed in both instances. The only experimental difference is that in Late Perception subjects aimed at their comfortable step length (recorded during baseline walking) while wearing the VR headset, whereas in Early Post-Adaptation they walked unconstrained (i.e., without any specific instruction) and without VR headset. Thus, washout of motor after-effects by the active perceptual task is expected, as indicated by the qualitatively smaller motor after-effects during Early Post-Adaptation compared to those during the Catch trial (colored dotted lines in Figure 3b). Two-way repeated measures ANOVAs (factors: epoch and speed) were used for each step length to show that motor after-effects are present after perceptual after-effects are washed out and that the motor after-effects are influenced by speed. Specifically, the slow and fast leg had significant resurgence of the motor after-effects following perceptual testing (Figure 3b: slow step length: p_epoch_<0.001; fast step length: p_epoch_<0.001) and the magnitudes of after-effects were regulated by walking speed (Figure 3b: slow step length: p_speed_=0.04, p_speed#epoch_=0.018; fast step length: p_speed_=0.041, p_speed#epoch_ =0.43). In summary, active perception after-effects decayed as subjects walked, but this did not fully washout motor after-effects.

**Figure 3.**
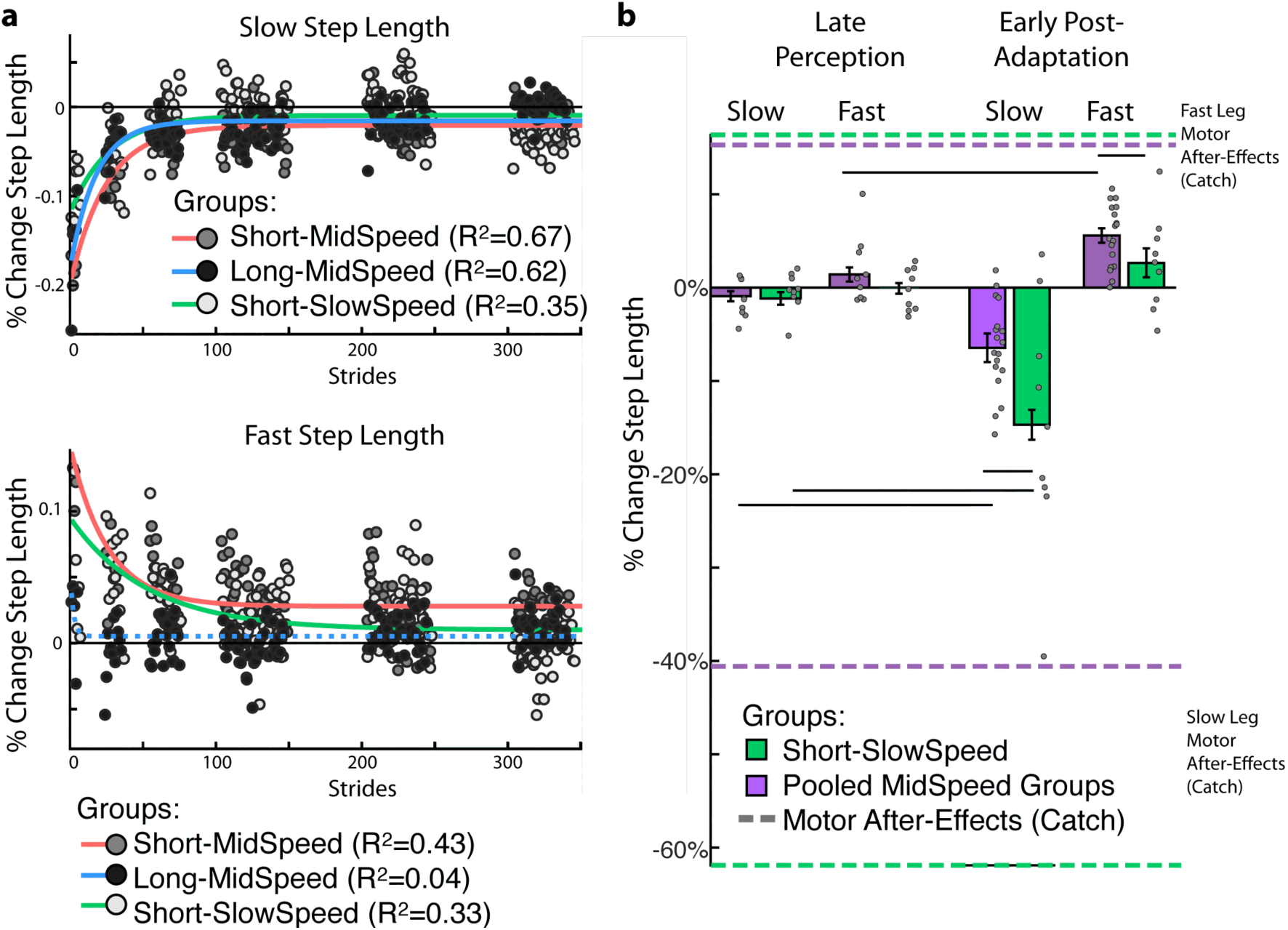
Active perception after-effects, but not motor after-effects, are fully washed out by the end of the perceptual task. **(a) Decay of active perception after-effects**. We show the %Change in Step Length (i.e., after-effects) for the slow (Top panel) and fast step length (Bottom Panel) over the course of the active perceptual task. Note that all groups exhibit an exponential decay of active perception after-effects in both legs except for the fast leg in the Long-MidSpeed group whose τ was not significantly different from zero (lack of significance indicated with a dotted line). This further indicates that subjects did not have significant fast step length perceptual shifts when probed with long step lengths. Perceptual after-effects assessed at target step lengths that were different from the initial probe were not included in the analysis (white spaces). **(b) Remaining motor after-effects following the active perceptual task.** Colored lines indicate the motor after-effects recorded during the Catch trial (prior to the active perceptual task, Figure 2b). Bars indicate group means for the slow and fast %Change Step Length at the end of the perceptual task (Late Perception) and at the beginning of the Post-Adaptation period (Early Post-Adaptation). Black lines indicate Fisher’s LSD post Hoc results. Subjects walked at the exact same tied-belt speeds during the Late Perception and Early Post-Adaptation periods. The only difference in these time points was that subjects were prompted to take their comfortable step length during Late Perception, but during Early Post-Adaptation subjects walked unconstrained (i.e., without any specific instruction of where to step). Standard errors are displayed in all plots. Note that the mid speed groups are pooled for this analysis since they experienced the same Adaptation protocol and target step lengths during Late Perception.

### Active Perception After-Effects are Mediated by Changes in Trailing Leg Perception

Step lengths during active perceptual trials were further decomposed into perceived leading (*α*) and trailing (X) leg positions (Figure 4a) to determine which leg’s position perception underlies changes in step length perception following split-belt adaptation. Our results show that only the trailing leg’s position (X) exhibited active perception after-effects (Figure 4b). These were quantified with symmetry measures of each leg’s X, such that values different from zero indicated active perception after-effects. We separately characterized *α* and X perceptual after-effects for short and long target step lengths with single exponential fits (*y* = *c* + *ae*^(−*x*/τ)^). A separate fit was performed for individuals walking slow vs. those walking at mid speeds. Fits of the data indicate that only the trailing leg position experiences large changes in perception that decayed as subjects experienced the active perceptual task. This is indicated by the decay of perceptual after-effects probed at either the short step length target (Figure 4b: τ [95% CI]; Short Target: MidSpeed= 12.5 [10.8, 14.9], SlowSpeed=36.9 [32.6, 42.7]) or long step length target (MidSpeed=37.5 [33.4, 42.8], SlowSpeed=41.0 [33.5, 52.9]). In contrast, the leading leg’s position exhibited perceptual after-effects that were very short lived, regardless of whether they were probed with the short step length target (Figure 4c: τ [95% CI]; Short Target: MidSpeed= 8.4 [6.4, 12.4], SlowSpeed=1.2 [-0.01, 2.4]) or the long step length target (MidSpeed=7.2 [4.7, 15.2], SlowSpeed=7.3 [3.8, 90.0]). Taken together, these results indicate that active perception shifts in step lengths are due to changes in the perception of the trailing leg’s position, and not the leading leg’s position.

**Figure 4.**
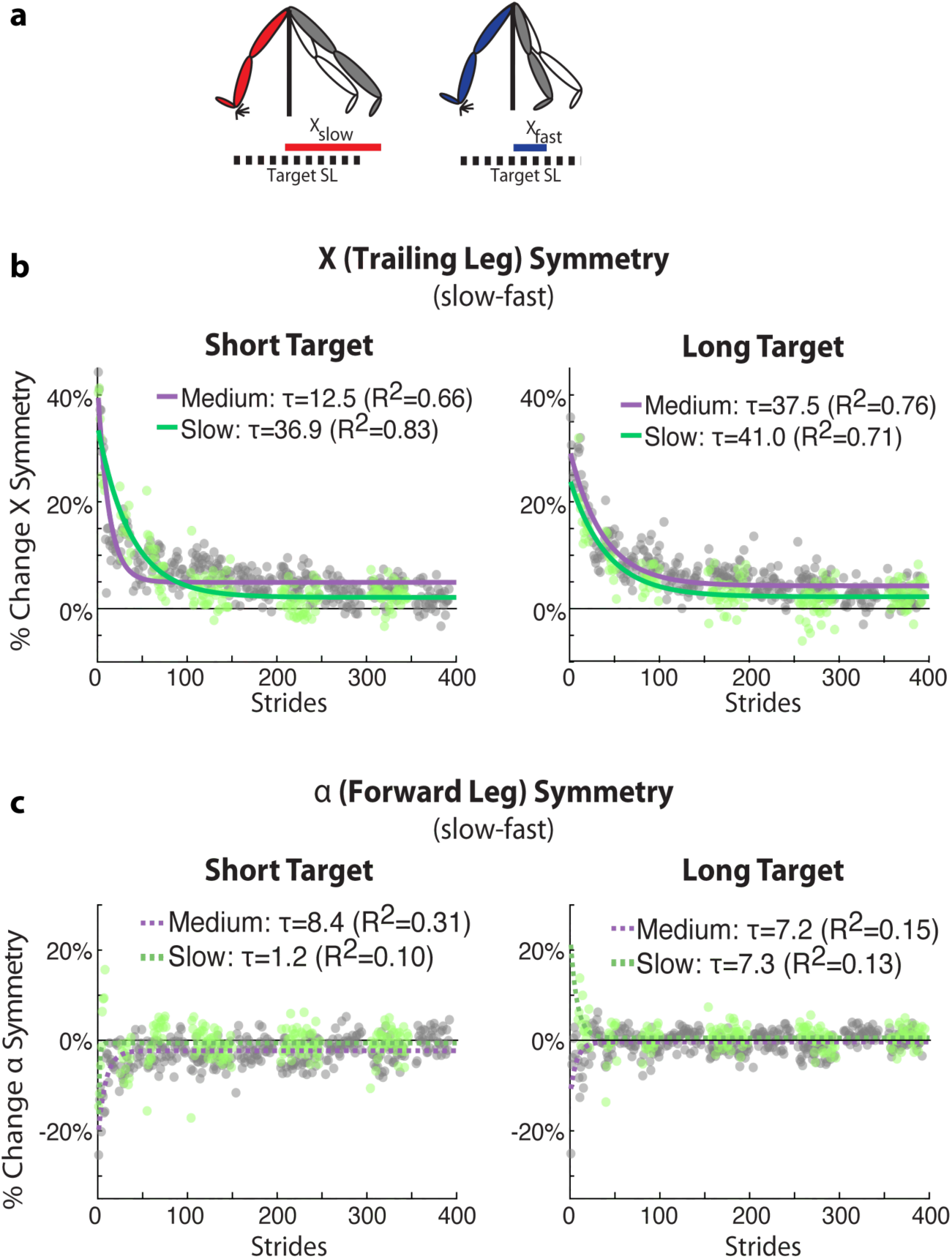
Split-belt Walking Induces Active Perception Changes in the Trailing Leg Position. **(a)** Schematic of the decomposition of step length into the leading (*α*) and trailing (X) leg relative to the hips. Both *α* and X are normalized by the speed specific baseline walking behavior, such that positive values mean the distance from the body is greater following Adaptation and vice versa. Symmetry measures were computed for both X **(b)** and *α* **(c)** and were defined as the differences between these measures on two consecutive steps (slow leg - fast leg). Thus, positive values of %Change X Symmetry here indicates that the slow leg (X_slow_) is trailing farther behind than fast leg (X_fast_) relative to baseline behavior. Similarly, positive values of %Change *α* Symmetry would indicate that the slow leg (*α*_slow_) is farther in front of the body than the fast leg (*α*_fast_) relative to baseline behavior. Note that the X position perception is altered following split-belt walking, but that the *α* position perception is not altered, as indicated by negligible %Changes in *α* Symmetry following split-belt walking.

## Discussion

### Summary

We investigated the effects of locomotor adaptation on passive and active step length perception. We observed active, but not passive, perceptual after-effects of step length. Thus, locomotor adaptation did not induce sensory shifts, but altered the estimation of limb position based on integration of afferent and updated efferent information. In addition, we found that active perception after-effects were only observed on the trailing limb’s position, indicating that the perceived position of the trailing, but not the leading leg, with respect to the body is recalibrated after locomotor adaptation. Moreover, perceptual and motor after-effects were regulated by how they were tested. Specifically, perceptual after-effects were altered by step length size, whereas motor after-effects were altered by walking speed. Lastly, the observed perceptual after-effects decayed as subjects walked in the active perceptual task, but motor after-effects remained, raising the possibility that after-effects in these two domains are originated by partially distinct processes.

### Motor recalibration in locomotion induces changes in active, but not passive perception of limb position

Our results show that the perception of limb position is altered following motor adaptation in locomotion, which supports prior findings that the perception of limb position is susceptible to motor adaptation (e.g., ^5,6,15,7–14^). However, we only observed perceptual after-effects when subjects actively generated motor commands and not when their legs were passively moved, which is consistent with prior work indicating that afferent signals encoding position remain intact following split-belt walking ^16^. These findings contrast multiple reports of shifts in the estimation of hand position following motor adaptation when the arms are moved by the experimenter (e.g., ^5–12^). We believe that this discrepancy between reaching and locomotion suggests that sensory changes post-adaptation arise from mismatched position estimates from different sensory sources (i.e., proprioception and vision), which is less prominent in walking. We also found that active perceptual effects were not observed in the leading leg’s position, as previously reported ^16^, but they were clearly observed in the trailing leg’s position, which had not been assessed before. We posit two potential explanations for the altered estimation of the trailing leg’s position. First, subjects might maintain the expectation that one leg will move faster than the other post-adaptation. As a result, they will stand longer on the leg that walked slow compared to the other as they take steps in the active perceptual task, leading to differences in trailing leg position that are consistent to the ones we see (i.e., X_slow_>X_fast_). Alternatively, the shifts in position might be due to changes in how subjects perceived the environment. It is known that the perception of symmetric walking speeds is shifted following split-belt walking ^16,25^. More specifically, individuals during post-adaptation perceive the fast leg to move slower and the slow leg to move faster than their actual speed. Subjects might act according to this perception by standing longer on their trailing leg when taking steps with the fast leg and vice versa for steps with the slow leg, resulting in the observed larger X_slow_ than X_fast_ values. In sum, subjects’ estimation of limb position is altered following sensorimotor adaptation. This shift is not due to changes to afferent information, but might be due to updated motor commands or altered perception of the environment following adaptation.

### Motor and perceptual after-effects are susceptible to how they are tested

It has been shown that split-belt motor after-effects are sensitive to walking speed ^20^ and perceptual after-effects are sensitive to movement distance ^18^. Thus, our observations that motor after-effects are regulated by walking speed and perceptual after-effects are regulated by step length are consistent with previous literature. However, it should be noted that walking speed and step length are not independent (e.g., ^22,23^) raising the possibility that step length is the factor that regulates both motor and perceptual after-effects. Should that be the case, we speculate that similarity in step lengths sizes during training and testing is a critical feature to maximize after-effects, given the well-known effect of similarity on the generalization of learned movements across conditions ^26–28^. In sum, our results highlight the importance of considering how both motor and perceptual after-effects are tested.

### Why did motor after-effects remain after perceptual ones were fully washed out?

It is interesting that walking in the active perceptual task did not fully eliminate motor after-effects. We believe this could be due to three possible reasons. First, the perceptual and motor effects were tested under different peripheral vision because subjects were wearing a virtual reality headset in the active perceptual task. This distinct situation constitutes a contextual difference known to reduce the washout of the pattern specific to the split-belt treadmill by walking in a symmetric environment ^29^. Second, the constrained nature of walking in the perceptual task might also influence the remaining motor after-effects. Namely, perceptual after-effects were tested by instructing subjects “where” to step, whereas motor after-effects were tested in unconstrained walking. Constrained and unconstrained movements are thought to be differently controlled (e.g. ^30^). Thus, the fact that individuals were aiming to a target in the perceptual task might have altered the extent to which implicit actions were washed out. Lastly, the limited washout of motor after-effects could also indicate that motor and perceptual after-effects are originated by partially distinct processes. This is supported by previous research showing that motor and perceptual effects exhibit different patterns of saving ^25^ and retention ^11^, distinct adaptation rates ^6,15^ and uncorrelated amounts of adaptation ^8,9,12,13^. They also exhibit different patterns of generalization to other limbs ^17^ or novel targets ^18,19^. Thus, while the limited washout of motor after-effects by the perceptual task might be due to testing differences, it could also indicate that the processes underlying motor and perceptual after-effects might be partially independent.

### Clinical implications

The split-belt walking may be a viable tool to address perceptual deficits associated with stroke. Notably, stroke survivors’ inability to perceive that their gait is asymmetric is associated with limited recovery of symmetric gait ^21^. Our results indicate that active perception of step lengths can change following split-belt walking, suggesting that a similar effect might be observed post-stroke. It has been shown that split-belt training can correct motor deficits ^31,32^ and it would be interesting to determine if it also corrects perceptual deficits as well. Of note, recent work indicates that perceptual changes in the estimation of speed are reduced with repeated exposure to the split condition ^25^, suggesting that similar reduction would be observed in the shifts of limb position. Thus, there might be a limited long term effect of split-belt walking on perceptual changes. Therefore, future work is needed to understand if split-belt walking has similar effects in the lesioned motor system and whether training protocols could lead to long lasting improvements in motor, as well as, perceptual deficits post-stroke.

## Methods

The influence of split-belt walking on passive (Experiment 1: n=8, 2 females, 24.8±4.8 yrs.) and active (Experiment 2: n=27, 16 females, 25.1±5.4 yrs.) step length perception was characterized in separate groups of subjects. This study was approved by the University of Pittsburgh Institutional Review Board, was carried out in accordance with the approved guidelines, and all subjects provided written informed consent.

### Experiment 1: Passive Perception

We tested subjects’ perception of their limb position by moving their legs in order to determine if afferent information was altered by split-belt walking. This was done before and after a tied and a split session, which occurred on different days. Both the tied and split sessions included perceptual and locomotor epochs (Figure 5a).

**Figure 5.**
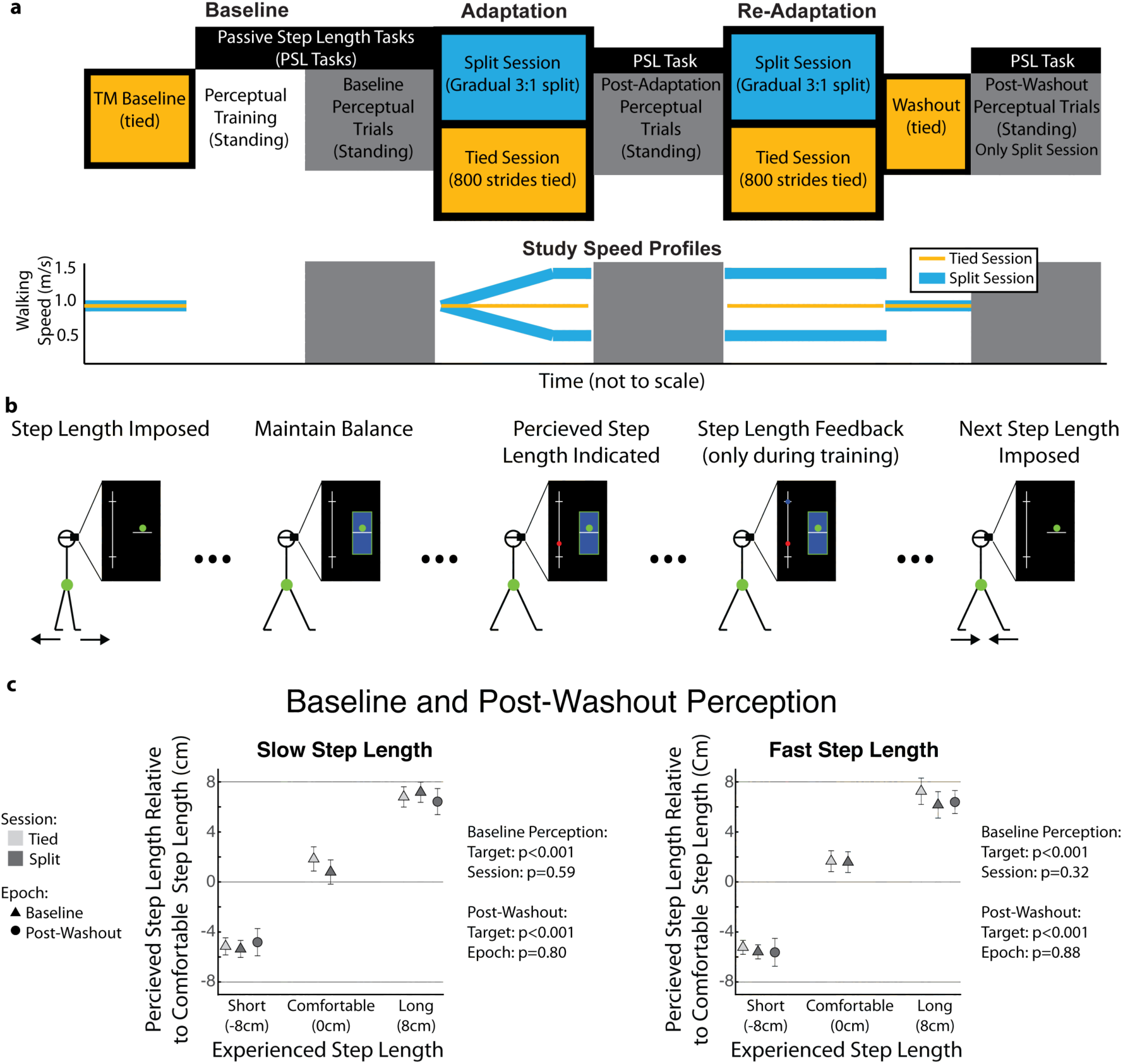
Locomotor and Passive Perception Protocol. **(a)** Locomotor epochs (colored blocks) and Passive Perceptual epochs (grey scale blocks) were included in both the control and testing session. Note that subjects were standing during the passive perceptual trials and walking during the locomotor trials. **(b) Passive Step Length Tasks:** Subjects’ legs were passively moved to a target step length. Subjects maintain an equal distribution of their weight between their feet using visual feedback displayed in the virtual reality headset. Subjects indicated their perceived step length size using an analog scale that was also displayed. Only during training, participants received feedback on their perceived vs. actual step length along the analog scale. This was done such that individuals could learn the mapping between what they felt and what was displayed in the analog scale. **(c) Baseline and Post-Washout Perceptual Trials:** Subjects’ performance during the Baseline Perceptual Trials for both sessions and the Post-Washout Trials for the split session are shown for each leg. Note that subjects could discern three distinct step lengths from the beginning (Baseline) until the end (Post-Washout) of the experiment. Thus, any performance changes between the Baseline vs. Post-Adaptation epochs were attributed to shifts in subjects’ passive perception of step length, rather than subjects’ inability to maintain the mapping between the imposed step lengths and the analog scale.

#### Locomotor Task

Subjects experienced a tied and split session that differed only in the locomotor Adaptation and Re-Adaptation epochs (Figure 5a). The first session was used to determine if an extended period of tied belt walking (i.e., walking with both belts moving 1.0 m/s) during the Adaptation epoch changed subjects’ passive perception of step length. During the second session, subjects walked with the legs moving at different speeds during the Adaptation epoch to determine if locomotor adaptation altered subjects’ passive perception of step length. The split-belt perturbation was gradually introduced during the Adaptation epoch for consistency with prior work studying passive perception (e.g., ^6,8,9,12,33^). More specifically, belts started at 1.0 m/s and linearly diverged for 600 strides until a 3:1 split-belt ratio (i.e., 1.5 m/s: 0.5 m/s) was achieved and maintained for 200 strides resulting in a total of 800 strides for the Adaptation epoch. Subjects’ passive perception of step length was assessed immediately before and after the Adaptation epoch. We re-exposed individuals to either the tied condition (i.e., both legs at 1 m/s) or split condition (i.e., 1.5 m/s: 0.5 m/s) during a Re-Adaptation epoch to subsequently evaluate motor after-effects in each subject. We chose to evaluate motor after-effects following the perceptual task to avoid changes in perception due to multiple transitions between tied and split environments, which are known to alter perceptual effects ^25^. The Re-Adaptation epoch lasted 300 strides to ensure individuals reached a similar steady state as in the Adaptation epoch prior to assessing motor after-effects. Then, we evaluated motor after-effects during a Post-Adaptation epoch (450 strides) when the two legs moved at the same speed (1 m/s). Subjects walked in this tied environment for 450 strides to fully wash out any split-belt after-effects before testing subjects’ retention of the mapping between the imposed step lengths and the analog scale used in the perceptual task (see section below).

#### Passive Perceptual Tasks

During the passive perception task, subjects wore a virtual reality headset (Oculus, Facebook Technologies, Irvine, CA, United States) while standing with their weight evenly distributed between the two feet and with each foot placed on one belt of the split-belt treadmill. The sequence of a passive perceptual trial is shown in Figure 5b. First, legs were passively moved to a specific step length value (ankle to ankle distance) (Figure 5b, first panel). Then, belts were stopped and subjects were asked to distribute their weight equally between their feet by maintaining their center of pressure (green dot) within a target region (blue rectangle) (Figure 5b, second panel). Subjects maintained this balanced position for 5 seconds during which they had to indicate their perceived step length using a clicker that moved a cursor (red dot) along an analog scale displayed in the virtual reality headset (Figure 5b, third panel). Then, a new perceptual trial was initiated by moving the legs to a different step length value (Figure 5b, fifth panel). Subjects received feedback of the actual step length in the analog scale (dark green dot) only during perceptual training (see full description below) (Figure 5b, fourth panel). Subjects did not lift their feet off the belts when the belts were moving or still. We alternated the leg that was placed forward after every perceptual probe/trial.

##### Perceptual Training

Subjects trained extensively on the passive step length task to learn the mapping between the imposed step lengths and the analog scale. Training occurred at the beginning of every session. Three out of the 8 participants had an additional session with only training trials prior to the tied session to determine if improved accuracy at the task would allow us to identify small perceptual shifts. The analog display observed by all subjects is shown in Figure 5b The middle of the analog scale corresponded to a medium size step length computed as 75% of each subject’s averaged step length during baseline walking at 1.0 m/s. Subjects additionally trained on targets 8 cm shorter (short step length) and longer (long step length) than the medium size step length. Six out of the eight participants on this task also trained with step lengths that were 4 cm shorter and longer than the medium size step length. During training, subjects experienced the same perceptual probe sequence as during testing: 1) legs were passively moved to one of the target step length values (e.g., medium, short, and long step length sizes), 2) subjects indicated their perceived step length while maintaining an equal distribution of their weight between their feet, and 3) subjects received feedback on their actual step length along the same analog scale. Subjects experienced 10 or 20 iterations of each of the target step lengths in a random order. Our results did not change when either more step length sizes were assessed (e.g., 3, rather than 5, target step lengths) or more instances of each step length size were tested (e.g., 21, rather than 10).

##### Perceptual Trials

We evaluated subjects’ capacity to classify step length sizes with the analog scale immediately following training (i.e., Baseline Perceptual epoch) and right after the Adaptation epoch during each session (i.e., Post-Adaptation epoch). Each epoch consisted of several iterations (10 or 21) of the target step sizes presented in a random order. Subjects performance was computed as the mean value that they selected in the analog scale across all perceptual trials at each target size. In all trials, subjects were told that they would experience step lengths all along the analog scale, including values that were shorter or longer than those they experienced during training. In reality, subjects were only exposed to the step length values that they trained on. Feedback was not provided in any trial. The performance of the perceptual trials for each leg were used to determine if subjects could differentiate the distinct step length sizes with the analog scale (i.e., Baseline Perceptual epoch performance), and if passive perception of step length was altered by the Adaptation epoch (Baseline Perceptual vs. Post-Adaptation epoch performance). First, the Baseline Perceptual performance for each leg was compared across all target sizes and sessions to determine if subjects could differentiate the distinct step length sizes with the analog scale for each session. Figure 5c indicates that subjects learned to discern the three step length values they experienced when their legs were passively moved (Figure 5c: two-way repeated measures ANOVA testing the effect of target and session on perceived step length; Slow Leg: p_target_<0.001, p_session_=0.59, p_target#session_=0.56; Fast Leg: p_target_<0.001, p_session_=0.32, p_target#session_ =0.71). Next, it was determined if passive perception of step length size was altered by split belt walking by comparing the Baseline Perceptual performance and Post-Adaptation performance (see results). Lastly, we probed subjects’ passive perception at the very end of the split session in a Post-Washout epoch, during which subjects’ perception of the short and long step length were probed 7 or 10 times each. This was done to ensure that changes in the reported step length with the analog scale were due to perceptual changes, rather than forgetting of the mapping between the sensed step length and the analog scale. Thus, subjects’ perception of the short and long step lengths were compared between the Baseline Perceptual and Post-Washout epochs. We found no differences between epochs or target step lengths (Figure 5c: two-way repeated measures ANOVA testing the effect of epoch and target on perceived step lengths: Slow Leg: p_epoch_=0.80, p_target_<0.001, p_epoch#target_=0.13; Fast Leg: p_epoch_=0.88, p_target_<0.001, p_epoch#target_=0.83). This indicated that individuals could classify well the distinct step length values throughout the duration of the experiment.

### Experiment 2: Active Perception

Subjects’ estimation of step length upon active motion of their legs was evaluated to determine if this estimation was altered by split-belt walking. This was done before and after a tied and a split session, which occurred on different days. Both the tied and split sessions included perceptual and locomotor epochs (Figure 6a). Importantly, perceptual and motor after-effects were tested under distinct walking speeds (i.e., mid and slow) since this factor has been shown to modulate the generalization of sensorimotor recalibration across walking conditions ^20^. Moreover, speed naturally regulates the magnitude of step lengths ^22,23^. Therefore, we evaluated the possibility that active perception of step length would be distinct for different step length sizes (i.e., long and short). Of note, both step length sizes were tested in all groups, but we altered the order at which the perception of step lengths were probed (i.e., long first vs. short first) given our expectation that active perceptual after-effects would decay as subjects performed the active perceptual task. In sum, motor and active perceptual after-effects were tested at either a slow (0.5 m/s) or mid (1 m/s) walking speeds, and active perceptual after-effects were first measured at either short or long step lengths. These experimental conditions resulted in three groups designated first by testing speed and second by the first step length size at which active perception was probed (Long-MidSpeed, Short-MidSpeed, Short-SlowSpeed).

**Figure 6.**
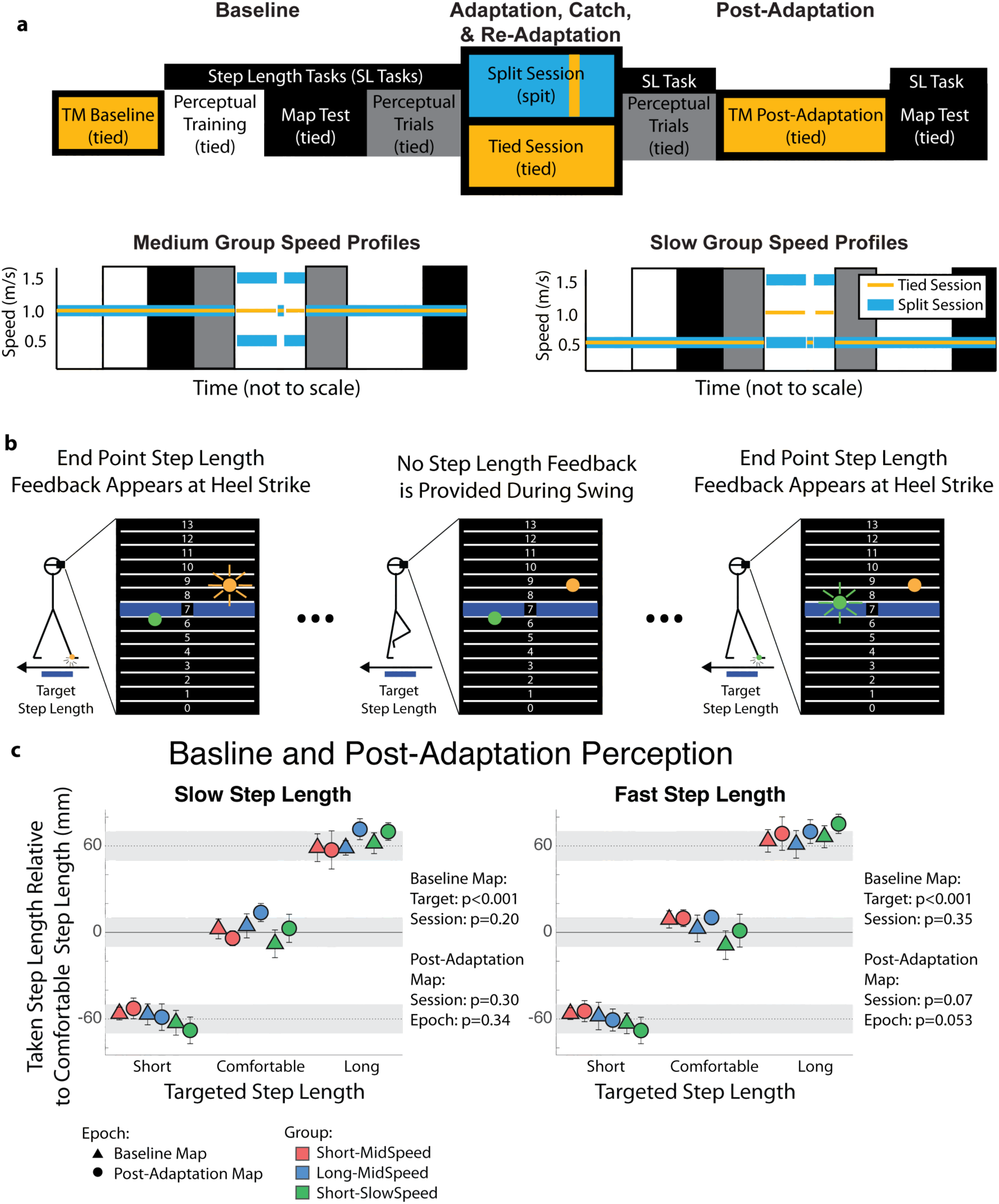
Locomotor and Active Perception Protocol. **(a)** Locomotor (colored blocks) and Active Perceptual (grey scale blocks) epochs were included in both the control and testing session. Note that subjects were walking during the active perceptual trials. **(b) Active Step Length Task:** Subjects walked on the treadmill while wearing a virtual reality headset. Note that subject’s walking speed depended on the group. Subjects saw a grid where each grid corresponded to 2 cm in real space. The blue highlighted grid line indicated a target step length that subjects should take. Grid 7, 10, and 13 corresponded to short, medium, and long step lengths, respectively. During training trials, subjects received accurate endpoint feedback on each step length. During Map Tests, subjects did not receive any feedback on their performance. During perceptual trials, subjects received reduced error feedback (i.e., subjects only saw 35% of the error that they made). **(c) Map Test:** Subjects performance during the Map Test indicated how well subjects learned and retained the spatial mapping between step lengths prompted with biofeedback and taken step lengths. Note that subjects learned distinct step lengths that were well maintained from the beginning until the end of the experiment. Thus, any changes in the active perception of step length are due to perceptual changes rather than subject’s inability to learn or maintain a mapping between prompted and taken step lengths.

#### Locomotor Task

Similar to Experiment 1, subjects experienced a tied and split session that differed only in the locomotor Adaptation and Re-Adaptation epochs (Figure 6a). The first session ensured that tied walking (i.e., walking with both belts moving 1.0 m/s) during the Adaptation and Re-Adaptation epochs did not induce active changes in step length perception. During the second session, subjects walked for 600 strides at a 3:1 split-belt perturbation (1.5 m/s and 0.5 m/s) during the Adaptation epoch. Unlike Experiment 1, we introduce the split-belt perturbation abruptly because it has been shown that active perception adapts more slowly than motor recalibration ^15^. Thus, we wanted to maximize the exposure of the full split condition before assessing perceptual after-effects. Motor after-effects were characterized in a short tied Catch trial at the group-specific walking speed. During the split session’s Re-Adaptation epoch, subjects again experienced 3:1 split-belt walking for 300 strides in order to return subjects to steady state split-belt walking before testing active perception of step length in the Post-Adaptation epoch. Following the active perceptual assessment during the Post-Adaptation epoch, subjects walked for 150 strides at the group-specific walking speed in order to evaluate any remaining motor after-effects. Once these motor after-effects were fully washed out, we performed one last active perceptual task without feedback to assess subjects’ retention of the map between the target and executed step lengths.

#### General Approach for all Active Perception Trials

Subjects wore a virtual reality headset (Oculus, Facebook Technologies, Irvine, CA, United States) to eliminate the possibility of using visual information to bias subjects’ estimate of step length based on their body motion with respect to the treadmill. Subjects received limited feedback on their step length on a numbered grid that was displayed in the virtual reality headset (Figure 6). Each grid line corresponded to 2 cm in real space. The cursors on the screen indicated the step length when taking a step with the right (orange cursor) or left leg (green cursor). The cursor position for each leg was only updated at heel-strike. The target step length was indicated with the blue highlighted line on the grid. The comfortable step length (grid #10) corresponded to the average step length during the speed-specific baseline walking. Subjects were also prompted to take a short (grid #7, 6 cm shorter than comfortable step length) and long (grid #13, 6 cm longer than comfortable step length) target step lengths. Importantly, the short and long step lengths used in the perceptual task were different from those regularly taken during baseline walking or during the catch trail. This was done to test active perceptual effects with step lengths that were distinct from those that subjects would normally take when assessing motor after-effects. Subjects performed the active perceptual task as they walked while lightly touching (<2N, enforced with verbal feedback) an instrumented handrail located in front of the treadmill. This was done to ensure that individuals would maintain their position on the treadmill while walking with the virtual reality headset ^34^. Recall that the Short-MidSpeed and Long-MidSpeed groups performed the step length tasks at a mid-walking speed (1.0 m/s), whereas the Short-SlowSpeed group performed the step length tasks at a slow walking speed (0.5 m/s).

##### Training for active perceptual testing

A step length Training epoch was first conducted in order to train subjects to take three distinct step lengths. During training, subjects received feedback on 100% of their step length error, where step length error is defined as the distance between the target step length and the executed step length. Subjects experienced targets in 4 sets where each set included each of the three targets in a random order. Each presentation of the target included 50 strides with visual feedback followed by 10 strides with no visual feedback. This was done to habituate the subjects to performing the perceptual task without feedback and favor the learning and retention process (e.g., ^35^). In total, subjects trained for 720 strides.

##### Map Test

Map Tests were used to ensure that subjects had successfully learned and maintained a mapping between the target step lengths prompted by each number on the grid. During this map test, subjects performed two sets of 25 strides for each target (total of 150 strides) without any feedback. This was done right before the Adaptation epoch and at the end of each session following the extinction of any locomotor after-effects during the Post-Adaptation epoch. The perceptual training protocol was effective as indicated by the fact that subjects took distinct step lengths for each target step length value (Figure 6c: two-way repeated measures ANOVA testing the effect of target step length and session on step length for each leg; Slow Leg: p_Target_<0.001, p_Session_=0.20, p_Target#Session_=1.00; Fast Leg: p_Target_<0.001, p_Session_=0.35, p_Target#Session_=0.80) that were retained for the duration of each session except for the fast leg during the tied session (two-way repeated measures ANOVA with effect of target step length and epoch; Tied Session, slow step length: p_Target_=0.82, p_Epoch_=0.79, p_Epoch#Target_=0.68; Tied Session, fast step length: p_Target_=0.92, p_Epoch_=0.009, p_Epoch#Target_=0.37; Split Session, slow step length: p_Target_=0.89, p_Epoch_=0.069, p_Epoch#Target_=0.80; Split Session, fast step length: p_Target_=0.76, p_Epoch_=0.13, p_Epoch#Target_=0.72). While the mapping between prompted and taken step lengths was not well maintained for the fast leg during the tied session, qualitatively, subjects still maintained distinct step lengths. Therefore, any changes observed between the Baseline and Post-Adaptation Perceptual trials are due to changes in perception, rather than subjects forgetting of the step length value that they were required to take when prompted by each number on the grid.

##### Active Perception Trials

To determine if active perception was altered, subjects needed to be able to have large step length errors without receiving feedback that would illicit strategic movement corrections that could override perceptual shifts. However, pilot data indicated that subjects needed some feedback to stay on task for the duration of the perceptual trials. Therefore, 35% of subjects step length error was projected during the active perceptual trials directly before and after the Adaptation epoch without the subjects’ knowledge in order to observe active perceptual shifts and their natural decay (e.g., if subjects had a 10 cm step length error, the feedback only indicated that there was a 3.5 cm step length error). Recall that the Short-MidSpeed and Short-SlowSpeed group started with short targets whereas the Long-MidSpeed group started with long targets. Subjects alternated between the short and long target step lengths in a predefined manner in order to characterize the decay of any temporally persistent perceptual shifts (455 strides total: 10 strides first target, 15 strides of the second target, one set of the short and long target for 15 strides, one set of the short and long target for 25 strides, three sets of the short and long target for 50 strides, 50 strides of the comfortable step length). Note that because all groups took the same number of strides, that it took the group walking slow (0.5 m/s) twice as long to complete the perceptual trials as the groups walking at the mid speed (1 m/s).

### Data collection

Kinematic data were collected to characterize subjects’ locomotor behaviors while walking on the treadmill during locomotor and perceptual trials. A motion analysis system (Vicon Motion Systems, Oxford, UK) was used to collect kinematic data at 100 Hz. A quintic spline interpolation was used to fill gaps in the raw kinematic data (Woltring; Vicon Nexus Software, Oxford, UK). Subjects’ movements were tracked via passive reflective markers placed bilaterally over the hip (greater trochanter) and ankle (lateral malleoulous) and asymmetrically on the thigh and shank to distinguish the legs. The duration of treadmill trials was defined by real time kinetic detection of heel strikes. Heel strikes were identified with raw vertical kinetic data collected from the instrumented treadmill (Bertec, Columbus, OH, United States).

#### Perceptual Task

Custom written Vizard (WoldViz, Santa Barbara, CA, United States) and MATLAB (The MathWorks, Inc., Natick, MA, United States) code was used to control the treadmill in order to impose desired step lengths during the passive perception tasks and to provide appropriate feedback during all perceptual tasks.

### Statistical Analysis

The statistical analyses were performed as reported in the text. A significance level of *α* = 0.05 was used for all analysis. Stata was used to perform all statistical analysis (StataCorp LP, College Station, TX).

## Author Contributions

G.T.-O. and C.S. were involved with the conception and design of the work. C.S. and M.G.-R. collected and analyzed the data. C.S. and G.T.-O. interpreted the results. C.S. drafted the manuscript, which was carefully revised by all authors. The final version of the manuscript has been approved by all the authors who agree to be accountable for all aspects of the work in ensuring that questions related to the accuracy or integrity of any part of the work are appropriately investigated and resolved. All authors qualify for authorship and all those who qualify for authorship are listed.

